# Loss of Adenomatous polyposis coli function renders intestinal epithelial cells resistant to the cytokine IL-22

**DOI:** 10.1101/479972

**Authors:** Yu Chen, Ian P. Newton, Maud Vandereyken, Ignacio Moraga, Inke Näthke, Mahima Swamy

## Abstract

Interleukin-22 (IL-22) is critical in maintaining homeostasis in the intestine by regulating the balance between pathogenic and commensal bacteria. IL-22 also promotes wound healing and tissue regeneration, which can support the growth of colorectal tumours. Mutations in the tumour suppressor Adenomatous Polyposis Coli gene (*Apc*) cause intestinal tumorigenesis and are a major driver of familial colorectal cancers. To understand the role of IL-22 in APC-mediated tumorigenesis, we analysed IL-22 signalling in wild-type (WT) and APC-mutant cells in murine small intestinal epithelial organoids and in mice. In WT epithelia, antimicrobial defence, mucus production, and cellular stress response pathways were most strongly upregulated by IL-22. Surprisingly, we found that although IL-22 activated STAT3 in APC-mutant cells, STAT3 target genes were not effectively induced. Our analyses revealed that *Apc*^*Min/Min*^ cells were resistant to IL-22 due to reduced expression of the IL-22 receptor, and increased expression of inhibitors of STAT3, including histone deacetylases. We further show that IL-22 induced expression of nitric oxide synthase in WT epithelial cells and corresponding DNA damage. These findings suggest that IL-22 does not promotes tumour formation by driving the proliferation of transformed intestinal epithelial cells. Rather, IL-22 increases genetic instability thereby accelerating transition from heterozygosity (*Apc*^*Min/+*^) to homozygosity (*Apc*^*Min/Min*^) to drive tumour progression.

## Introduction

Chronic inflammation is positively associated with intestinal tumorigenesis (Mager *et al*, 2016; Eaden *et al*, 2001; Mizoguchi *et al*, 2011). Recent data has particularly implicated the cytokine Interleukin 22 (IL-22) in colorectal cancers (CRC). Consistently, IL-22 producing cells are enriched in human CRC and associated with its development (Jiang et al. 2013; Huang et al. 2015). IL-22 is produced by type 3 innate lymphoid cells (ILCs), CD4^+^ and CD8^+^ T cells and γδ T cells during intestinal inflammation (Mizoguchi, 2012; Pickert *et al*, 2009; Kryczek *et al*, 2014; Jiang *et al*, 2013; Ahlfors *et al*, 2014). IL-22 binds to the IL-22 receptor, which is composed of two heterodimeric subunits, IL-22RA1 and IL-10R2 (Xie *et al*, 2000; Kotenko *et al*, 2001). In the intestine, IL-22 receptor expression is restricted to non-hematopoietic cells, particularly intestinal epithelial cells (IEC) (Nagalakshmi *et al*, 2004). IL-22 activates the STAT3 (signal transducer and activator of transcription 3) pathway and regulates intestinal epithelial functions including tissue repair, mucus secretion and antimicrobial activity (Zheng *et al*, 2008; Pickert *et al*, 2009; Tsai *et al*, 2017; Bel *et al*, 2017; Nagalakshmi *et al*, 2004; Lindemans *et al*, 2015; Turner *et al*, 2013). Increasing evidence indicates that IL-22 not only facilitates tissue-protection, but also has pro-survival and proliferative effects that can promote colorectal cancer (Lindemans *et al*, 2015; Huber *et al*, 2012; Kryczek *et al*, 2014; Kirchberger *et al*, 2013).

Inactivating mutations in the tumour suppressor Adenomatous polyposis coli gene (*APC*) cause intestinal tumorigenesis and are a major cause of hereditary colorectal cancer (Rowan *et al*, 2000; Nelson & Näthke, 2013), a condition known as Familial adenomatous polyposis (FAP). In FAP patients, adenomatous polyps are present throughout the colon and develop into cancer if left untreated. In addition to FAP, inactivating mutations in the *APC* gene are present in more than 80% of non-hereditary colorectal cancers (Kinzler & Vogelstein, 1996). APC is best known as a negative regulator of Wnt signalling, contributing to regulation of cell proliferation and differentiation (Dikovskaya *et al*, 2007; Nelson & Näthke, 2013; McCartney & Näthke, 2008; Stamos & Weis, 2013). The *Apc*^*Min/+*^ (Min, multiple intestinal neoplasia) mice mimic FAP intestinal tumorigenesis and carry a truncated version of the *Apc* gene on one allele. A spontaneous loss of heterozygosity (LOH) in intestinal epithelial cells leads to loss of the wild type (WT) *Apc* allele. The resulting increased Wnt signalling and other epithelial changes together lead to adenoma (polyp) formation in the intestine. In this and other *Apc*^−^ deficient mouse models, inflammatory cytokines are established augmenting factors for intestinal cancer development (Wang *et al*, 2014; Putoczki *et al*, 2013). Consistent with the idea that IL-22 might also play a role in this process, *Il22*^*-/-*^ *Apc*^*Min/+*^ mice develop smaller and fewer tumours than *Il22*^*+/+*^ *Apc*^*Min/+*^ mice (Huber *et al*, 2012). However, it is not clear how IL-22 impacts on APC-mutant cells to increase tumorigenesis.

The aim of our study was to understand how IL-22 contributes to intestinal tumorigenesis in *Apc*^*Min/+*^ mice and determine whether loss of APC function affects the cellular response to IL-22. To this end, we performed RNA sequencing on intestinal organoids from wild type (WT) and *Apc*^*Min/+*^ mice composed of *Apc*^*Min/Min*^ cells to measure intestinal epithelial responses to IL-22 stimulation. Organoids are primary IEC cultures that grow in three dimensions (Sato *et al*, 2009). Importantly, organoids contain only IEC including stem cells, Paneth cells, enteroendocrine cells, goblet cells and enterocytes, but no immune or stromal cells, allowing us to identify the direct effects of IL-22 on small intestinal epithelia. We found that, surprisingly, APC-deficient cells were resistant to stimulation by IL-22. This was due to both, reduced expression of the IL-22 receptor and high expression of negative regulatory Histone deacetylases. Conversely, *Apc* ^*Min/+*^ heterozygous cells responded to IL-22 similarly to wild type cells by upregulating cellular stress responses involved in the production of reactive nitrogen species. This stress response correlated with enhanced DNA damage, which in turn, correlated with the accelerated transformation of *Apc*^*Min/+*^ cells to *Apc*^*Min/Min*^ genotype in the presence of IL-22, that ultimately leads to development of adenomas.

## Results

### The IL-22 regulated transcriptome in small intestinal epithelial cells

IL-22 regulates multiple genes involved in antimicrobial defence, proliferation and immune signalling. However, this varies according to cell type (Zheng *et al*, 2008; Pham *et al*, 2014; Wolk *et al*, 2006), and effects of IL-22 on primary small intestinal (SI) epithelial cells are currently unknown. To identify the responses of small intestinal epithelial cells to IL-22, we derived primary organoid cultures from wild-type (WT) murine small intestines and treated them with IL-22. We performed deep RNA sequencing (RNAseq) on WT small intestinal organoids 3 hours after treatment with IL-22, to measure the global IL-22–regulated transcriptome. We chose this time point as a key IL-22 target gene and negative feedback regulator, *Socs3*, was maximally induced at this time point (Fig. 2C), and we were interested in the immediate early effects of IL-22. The results showed that IL-22 significantly affected expression of 437 genes in WT organoids (Fig. 1A, S1A). Overwhelmingly, most of these were involved in immune response and defence against infection and stress (Fig. 1B, C). These included genes with direct antimicrobial activity (e.g. *Reg3g, Lcn2*), genes involved in pathogen recognition and immune signaling (e.g. *Tmem173 (*STING), *myd88)*, genes involved in mucus production and mucin glycosylation (e.g. *Fut2, B4galt1*), and those involved in the production of reactive nitrogen and oxygen species (e.g. *Nos2, Duox1/2*) (Fig. 1B). Other upregulated genes were involved in wound repair (e.g. *Areg*), and have been recently implicated in colorectal cancer cell proliferation, such as *Steap4*, a metalloreductase (Xue *et al*, 2017) and *Lrg1 (*Zhang *et al*, 2016). Notably however, no cell cycle genes were induced by three hours, and many metabolic pathways were downregulated, including the Aryl hydrocarbon receptor pathway, recently implicated in maintaining the intestinal stem cell niche (Metidji *et al*, 2018), and cellular lipid metabolism. We also did not find upregulation of genes previously implicated in ‘cancer stemness’, such as Nanog, Sox2 or DOT1L (Kryczek *et al*, 2014). However, we did detect some overlap with genes that have also been found to be upregulated by IL-22 in colonic organoids (Pham *et al*, 2014). Notable exceptions were genes involved in retinoic acid metabolism and vasculature development, which were not enriched in SI organoids (Fig. S1C-D). In summary, the main role of IL-22 in the small intestine appears to be protection from infection, through synthesis of antimicrobial proteins, increased mucus production, and oxidative stress.

**Figure 1.**
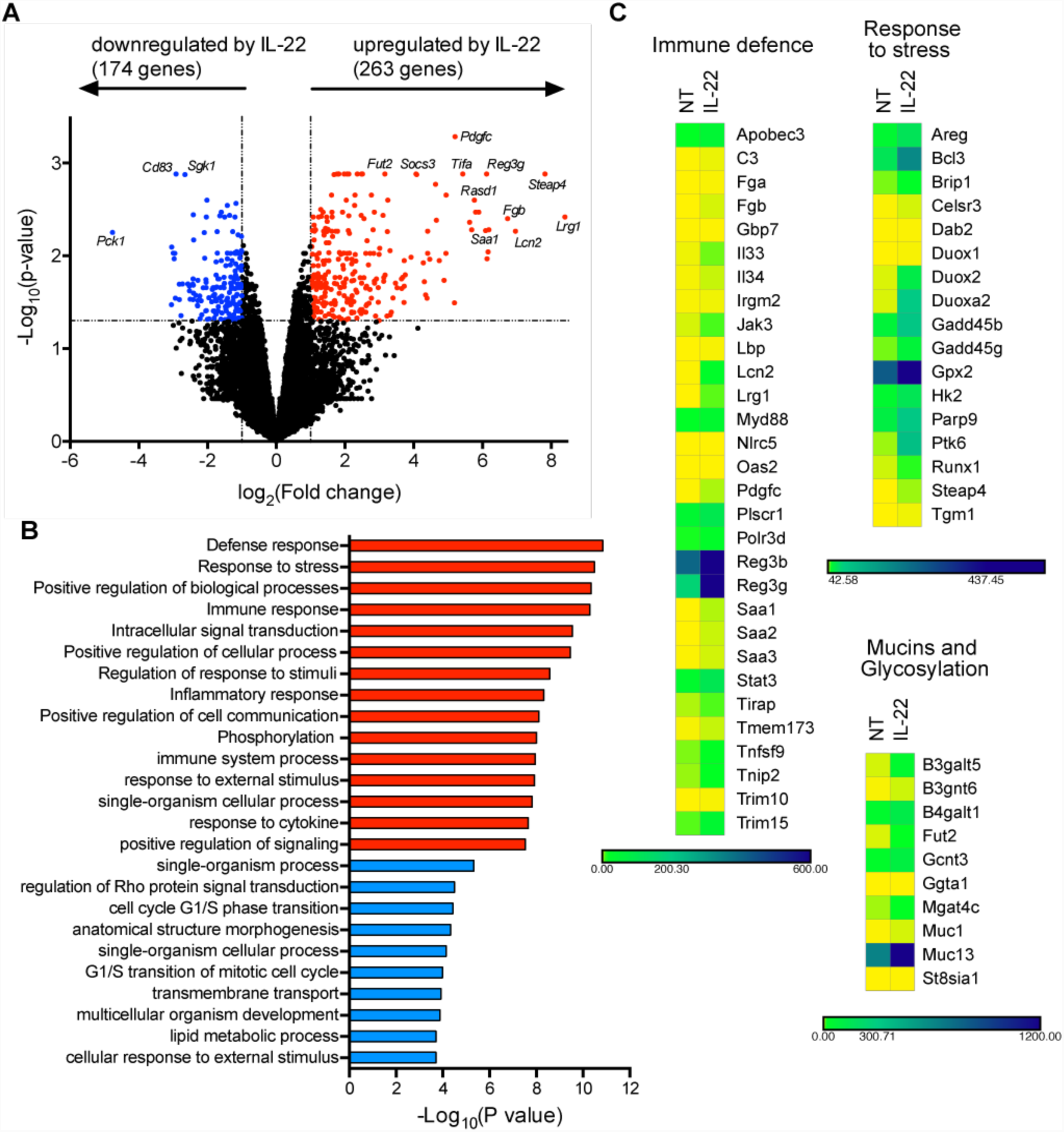
IL-22 regulates genes involved in intestinal immune defence responses. WT organoids were treated with IL-22 (2ng/ml) for 3 hours. RNA was isolated for RNA sequencing (RNAseq) analysis. (A) Volcano plot shows gene expression of WT organoids treated with IL22 (2ng/ml) for 3 hours (WT-IL22) compared to untreated WT organoids. Red=up-regulated gene (fold change > 2, p < 0.05), blue=down-regulated genes (fold change < 0.5, p < 0.05). (B) Functional enrichment analysis (GO term – Biological processes) for genes up-or downregulated in WT organoids treated with IL-22, compared to untreated WT organoids. Shown are the top 15 upregulated (red) and 10 downregulated pathways (blue). (C) Heat maps displaying mRNA expression level of significantly regulated genes in WT organoids either not treated (NT) or treated with IL-22. Genes of interest were manually grouped by their main biological function.

**Figure 2.**
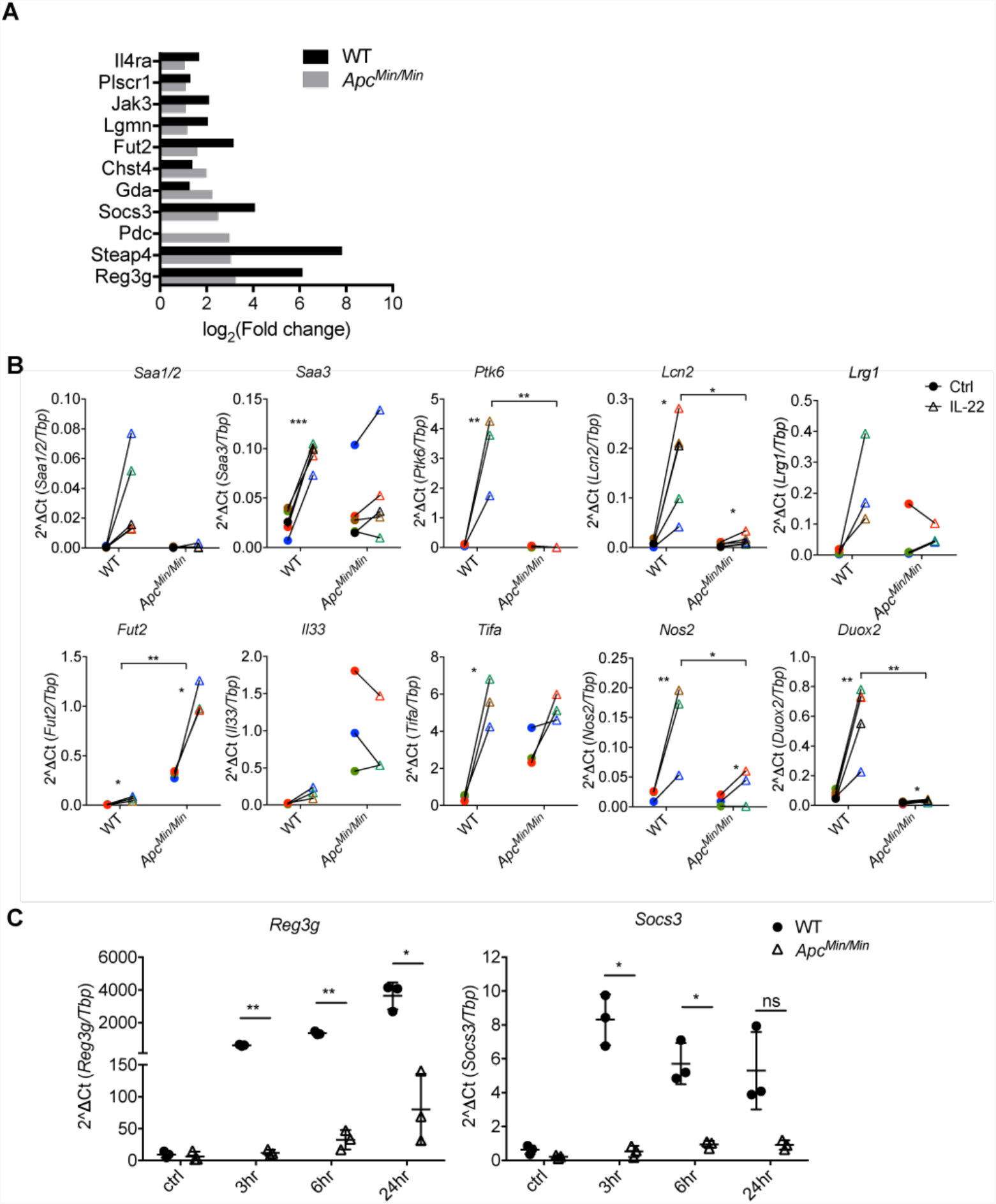
APC-mutant IECs are defective in IL-22 mediated gene regulation. (A). WT and *Apc*^*Min/Min*^ organoids were treated with IL-22 (10ng/ml) for 3 hours. RNA was isolated for RNAseq analysis. Graph shows the fold change of genes induced by IL-22 in WT and *Apc*^*Min/Min*^ organoids. The 11 genes shown were the only genes up-regulated by IL-22 in *Apc*^*Min/Min*^ organoids (p<0.1). (B). Expression of candidate up-regulated genes from RNAseq data were verified by RT-qPCR. Data shown are relative levels of mRNA for genes of interest relative to mRNA of TATA box binding protein (*Tbp*). At least 3 independent experiments were performed in each case. *P<0.05, **P<0.01, ***P<0.001 on paired t test. (C). WT and *Apc*^*Min/Min*^ organoids were treated with IL22 (10ng/ml) for 3, 6 or 24 hour and RT-qPCR analysis were performed. Data show the relative expression of mRNA for *Reg3g* or *Socs3* compared to *Tbp*. 3 independent experiments were performed. *P<0.05, **P<0.01, paired t test.

### Defective IL-22 mediated gene regulation in APC-mutant IEC

Since IL-22 has been implicated in promoting stemness and proliferation in colorectal cancer cell lines (Kryczek *et al*, 2014; Kirchberger *et al*, 2013; Lindemans *et al*, 2015), we asked how transformed SI epithelial cells responded to IL-22. We treated transformed *Apc*^*Min/Min*^ SI organoids (Langlands *et al*, 2018) derived from *Apc*^*Min/+*^ mice with IL-22, and performed RNASeq. In contrast to WT cells, we found only a minute number of genes regulated by IL-22 in APC-mutant IEC, and these were only detectable when reducing the stringency of the p-value threshold to p < 0.1 (Fig. S1A). 10 of the 11 upregulated genes were also induced by IL-22 in WT cells, indicating that IL-22 did not activate a different gene transcription programme in transformed cells (Fig. 2A). The non-responsiveness of *Apc*^*Min/Min*^ organoids to IL-22 was confirmed by quantitative reverse transcription-polymerase chain reaction (qRT-PCR) analysis (Fig 2B). Note that even at later time points, the gene transcription response to IL-22 was similarly reduced (Fig 2C) confirming that IL-22 did not activate a gene transcription program in *Apc*^*Min/Min*^ organoids.

### Transformed IEC respond poorly to IL-22 due to specific downregulation of the IL-22 receptor

To understand the almost complete lack of response to IL-22 in *Apc*^*Min/Min*^ organoids we investigated the activation of components of IL-22 signalling pathway. The most prominent effect of IL-22 in epithelial cells is activation of JAK1/Tyk2–mediated phosphorylation of STAT3 at Tyr705 to regulate gene expression (Dudakov *et al*, 2015). We first measured STAT3 expression by flow cytometry and confirmed that WT and *Apc*^*Min/Min*^ organoids have similar levels of STAT3 (Fig. S2A). We then stimulated organoids with IL-22 and measured STAT3 phosphorylation at the activating residue Tyr705 (pSTAT3) by immunostaining. IL-22 induced robust STAT3 phosphorylation in both WT and *Apc*^*Min/Min*^ organoids (Fig. 3A) with corresponding accumulation of pSTAT3 in the nucleus. However, consistently lower levels of pSTAT3 were induced in *Apc*^*Min/Min*^ organoids compared to WT organoids (∼50% lower, Fig. 3B). Moreover, this reduced response to IL-22 was neither improved after longer times nor when higher doses of IL-22 were used (Fig. 3C, D). Phos-tag gels confirmed that a smaller proportion of the total STAT3 was phosphorylated in response to IL-22 in *Apc*^*Min/Min*^ organoids (Fig. S2B).

**Figure 3.**
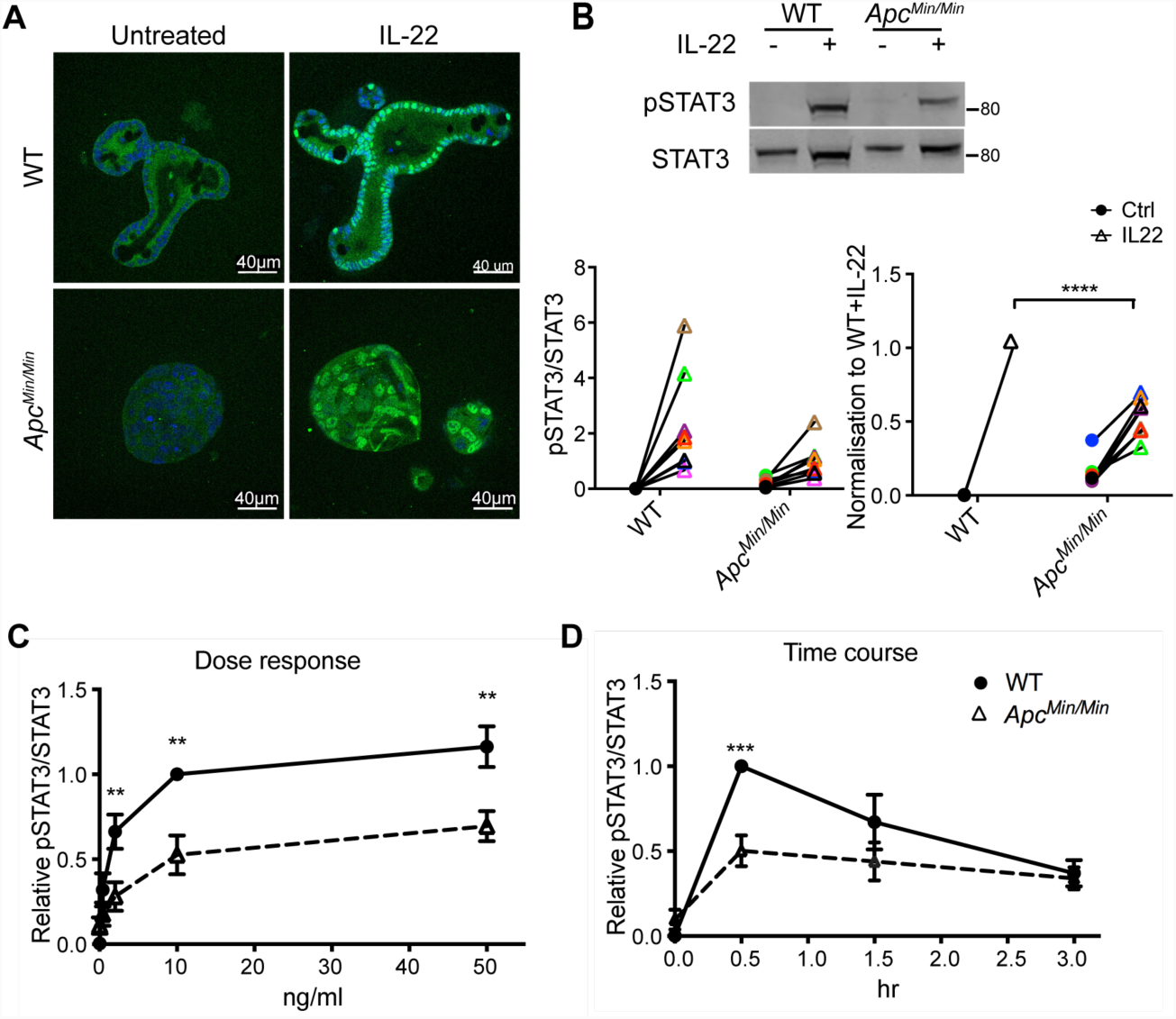
Differences in IL-22-induced STAT3 phosphorylation in WT and *Apc*^*Min/Min*^ organoids. (A). WT and *Apc*^*Min/Min*^ organoids were stimulated with IL22 (10ng/ml) for 0.5 hour. Organoids were fixed and immunostained with STAT3 antibodies (green) and nuclei were visualised with Hoechst (blue). (B). Immunoblots and corresponding quantification of pSTAT3 (Tyr705) in WT and *Apc*^*Min/Min*^ organoids with or without IL-22 stimulation. Relative level is expressed as the ratio of pSTAT3(Tyr705) to total STAT3 in each sample and normalised to the corresponding ratio in WT organoids treated with IL-22 in each experiment. (C). Dose response of pSTAT3(Tyr705) in organoids treated with 0.4, 2, 10 or 50 ng/ml IL-22 for 0.5 hours. (D). Time course of pSTAT3(Tyr705) in organoids stimulated with IL-22 (10ng/ml) for 0.5, 1 or 3 hours. pSTAT3/STAT3 levels in C and D were quantified by immunoblotting and normalised to WT + IL-22 at 10ng/ml IL-22 and at 0.5 hours respectively. *P<0.05, **P<0.01, ***P<0.001, two-way ANOVA.

These data indicated that transformed *Apc*^*Min/Min*^ IEC are intrinsically defective in activating Jak1/Tyk2 mediated phosphorylation of STAT3 in response to IL-22. To determine whether this effect was specific to IL-22, we tested whether other cytokines also produced reduced activation of STATs in *Apc*^*Min/Min*^ compared to WT organoids. IL-6 increases pSTAT3 in intestinal epithelial cells (Aden *et al*, 2016), and IFNα activates STAT1 phosphorylation (Giles *et al*, 2017). We treated organoids with the designer cytokine, hyper IL-6 (a fusion of the soluble IL-6 receptor (sIL-6R) with IL-6), which bypasses the requirement for the IL-6 receptor α chain to directly activate the gp130 receptor (Fischer *et al*, 1997), or with IFNα. We found that hyper IL-6 induced pSTAT3 (Fig. 4A) and IFNα induced pSTAT1 (Fig. 4B) similarly in WT and *Apc*^*Min/Min*^ organoids (Fig. S2D). Hence, the reduced levels of pSTAT3 in *Apc*^*Min/Min*^ organoids are specific to IL-22 responses.

**Figure 4.**
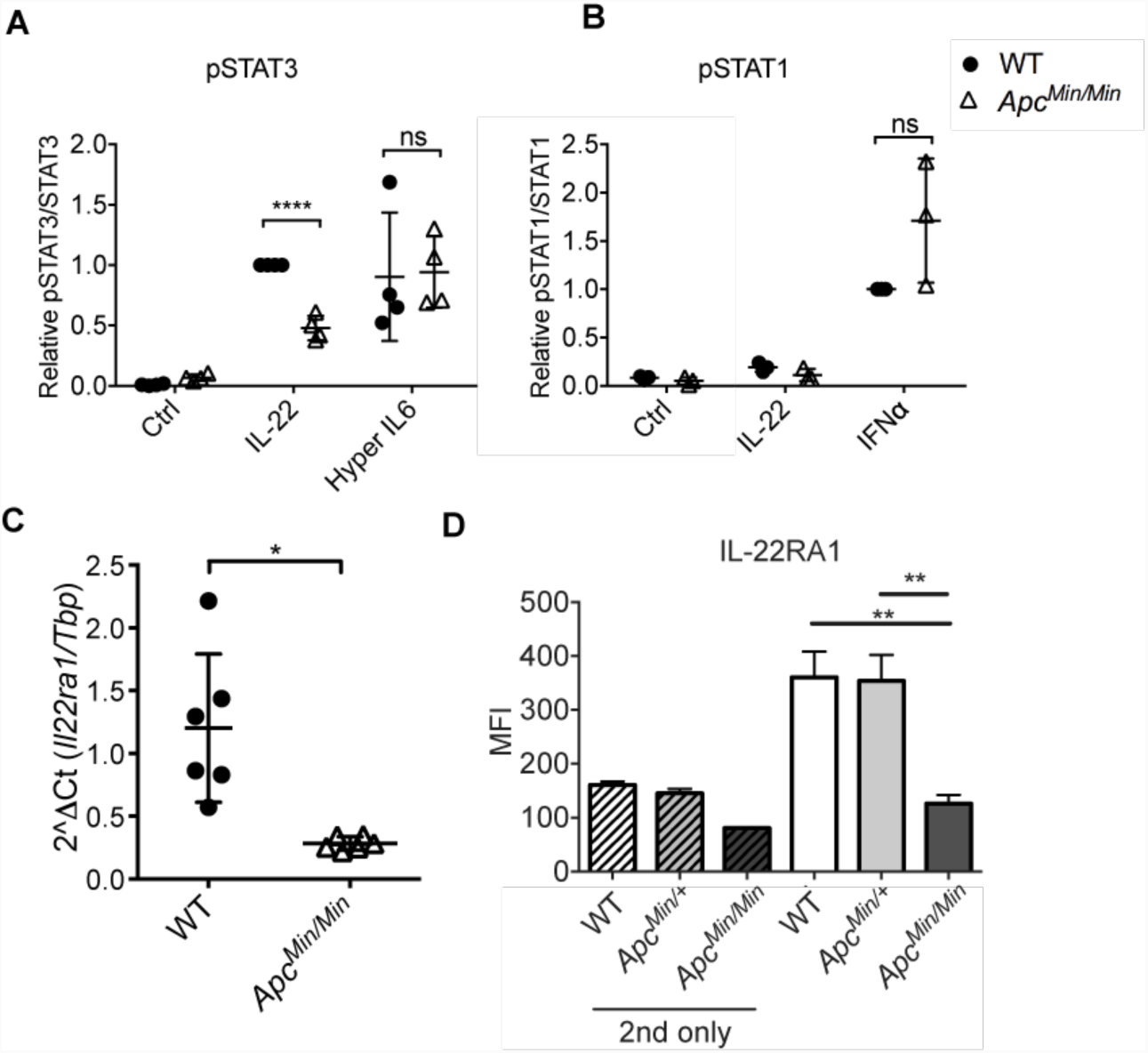
*Apc*^*Min/Min*^ organoids lose expression of the IL-22 receptor. (A) Amount of pSTAT3(Tyr705)/STAT3 in WT and *Apc*^*Min/Min*^ organoids stimulated with IL-22 (10ng/ml) or hyper IL-6 (50µM) for 0.5 hours, normalised to that in WT treated with IL-22 in each experiment. **P<0.01, paired t test. (B) pSTAT1(Tyr701) /STAT1 in WT and *Apc*^*Min/Min*^ organoids untreated or stimulated with IL-22 or IFNα (1000U/ml) for 0.5 hours, normalised to that in WT cells treated with IFNα in each experiment. (C) RT-qPCR analysis of *Il22ra1* expression in WT and *Apc*^*Min/Min*^ organoids. Six independent experiments were performed. *P<0.05, paired t test. (D) WT, *Apc* ^*Min/+*^ or *Apc* ^*Min/Min*^ organoids were dissociated and IL22RA1 expression was measured by flow cytometry. Mean fluorescence intensity (MFI) shows mean values ± SD of 3 biological replicates. ‘2nd only’ reflects samples that were stained only with 2^nd^ antibody (i.e. not exposed to primary antibodies). **P<0.01 on one-way ANOVA.

These results suggest that the reduction in pSTAT3 levels in *Apc*^*Min/Min*^ cells in response to IL-22 was caused by differences upstream of JAK/STAT activation, at the level of the cytokine receptor. Indeed, our RNAseq data revealed that mRNA for both IL-22 receptor subunits, *Il22ra1* and *Il10rb*, was lower in *Apc*^*Min/Min*^ than in WT organoids (Fig. S3A). Further analysis by RT-qPCR and flow cytometry confirmed that *Apc*^*Min/Min*^ IEC have significantly lower levels of *Il22ra1* mRNA with corresponding reduced IL-22 receptors on the cell surface (Fig. 4C, D). This could explain the lower induction of pSTAT3 specifically in response to IL-22 in APC-mutant cells, as gp130 expression (*il6st*) was not downregulated (Fig. S3A).

### Elevated levels of HDACs in APC-mutant IEC inhibit STAT3 target gene transcription

Hyper IL-6 induced pSTAT3 to similar levels in WT and *Apc*^*Min/Min*^ IEC. Nonetheless, induction of STAT3 target genes *Reg3g* and *Socs3*, was lower in *Apc*^*Min/Min*^ cells in response to IL-6 (Fig. 5A). On the other hand, IFNα-STAT1 target genes were equivalently induced in WT and *Apc*^*Min/Min*^ cells (Fig. 5B). This implied the transcriptional activity of STAT3, but not STAT1, was compromised in *Apc*^*Min/Min*^ IEC.

**Figure 5.**
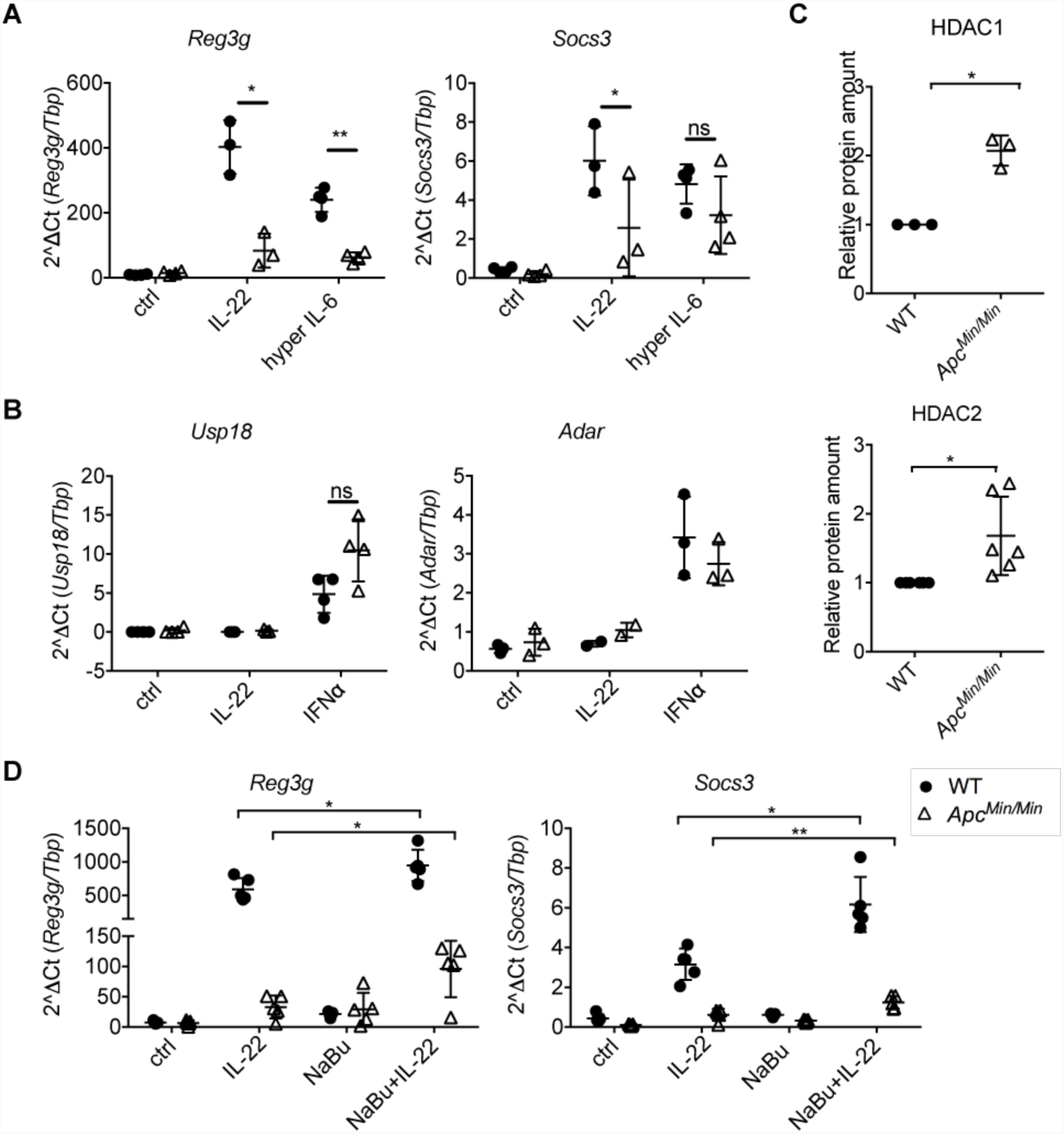
HDAC inhibition partially rescues STAT3-mediated gene transcription in *Apc*^*Min/Min*^ organoids. (A) RT-qPCR analysis of wildtype and *Apc*^*Min/Min*^ organoids treated with IL-22 (10ng/ml) or hyper IL-6 (50µM) for 3 hours. Data shows mRNA expression of STAT3 target genes, *Reg3g* or *Socs3*, relative to *Tbp*. (B) RT-qPCR analysis for WT and *Apc*^*Min/Min*^ organoids treated with IL-22 (10ng/ml) or IFNα (1000U/ml) for 3 hours. Data shows the expression of STAT1 target genes, *Adar* and *Usp18*, relative to *Tbp*. At least 3 independent experiments were performed. *P<0.05, **P<0.01, paired t test. (C) Western blot analysis of proteins in WT and *Apc*^*Min/Min*^ organoids. Expression of HDAC1 and HDAC2 was normalized to loading control, STAT3, in each sample and expressed relative to that in WT organoids. (D) WT and *Apc*^*Min/Min*^ organoids were treated with sodium butyrate (10mM) for 16 hours and then stimulated with IL22 (10ng/ml) for 3 hours. RT-qPCR was performed. Data show the levels of mRNA for *Reg3g* or *Socs3* relative to *Tbp*. *P<0.05, **P<0.01, paired t test. At least 3 independent experiments were performed.

Transcription by STAT3 is affected by a number of factors. In addition to phosphorylation at Tyr705, IL-22 increases the phosphorylation of STAT3 at Ser727, a site required for maximal STAT3 transcriptional activation (Wen *et al*, 1995). In contrast to pSTAT3(Y705) induction, which was impaired in *Apc*^*Min/Min*^ cells, we did not observe a difference in the level of IL-22 induced pSTAT3 (Ser727) in WT and *Apc*^*Min/Min*^ organoids (Fig. S2C). Once STAT3 is activated, it recruits coactivators such as Ncoa1 or CBP/p300 to the promoter of a target gene. In addition, its activity can also be negatively regulated by Type I histone deacetylases (HDAC), PIAS3 (protein inhibitor of activated STAT3), and SOCS3 (Icardi *et al*, 2012; Yuan *et al*, 2005; Liu *et al*, 1997; Giraud *et al*, 2002; Starr *et al*, 1997). Analysing our RNAseq data revealed lower gene expression of the STAT3 coactivator *Ncoa1* in APC-mutant cells, and higher expression of *Hdac1/2 (*Fig. S3B), but not of any of the other co-regulators. We also detected higher protein expression of HDAC1 and HDAC2 in *Apc*^*Min/Min*^ organoids (Fig. 5C). To further determine whether elevated HDACs contributed to lower expression of STAT3 target genes in APC mutant cells, we measured whether inhibition of HDACs by sodium butyrate (NaBu) could enhance STAT3 transcriptional activity. RT-qPCR results showed that pre-treatment of organoids with NaBu up-regulated IL-22 mediated STAT3 target genes, *Reg3g* and *Socs3*, in both WT and *Apc*^*Min/Min*^ organoids (Fig. 5D). Although *Reg3g* and *Socs3* gene expression was not rescued to WT levels in *Apc*^*Min/Min*^ cells, the relative increase in *Reg3g* after HDAC inhibition was higher in *Apc*^*Min/Min*^ cells compared to WT cells, reflecting higher HDAC expression. Thus, the defective response of transformed IEC to IL-22 is potentially a combined effect of both reduced expression of the IL-22 receptor and the high levels of HDACs expressed in *Apc*^*Min/Min*^ cells.

### Adenomas *in vivo* are also defective in responding to IL-22

Our observations so far focused on IEC in organoids. To test whether the differential response to IL-22 also existed *in vivo*, we injected WT or *Apc*^*Min/+*^ mice with IL-22 and measured STAT3 phosphorylation. In WT intestinal epithelia, pSTAT3 expression was not observed in PBS-injected control. Upon IL-22 treatment, nuclear pSTAT3 was strongly induced in WT intestinal tissue. In *Apc*^*Min/+*^ mice, IL-22 also induced pSTAT3 nuclear accumulation in untransformed cells. In polyps, which are comprised of homozygous *Apc*^*Min/Min*^ cells, a weak signal for nuclear pSTAT3 was already detectable prior to IL-22 treatment. However, there was no significant increase of pSTAT3 in polyps after injecting *Apc*^*Min/+*^ mice with IL-22 (Fig. 6A). Note that *Apc*^*Min/Min*^ tissue/polyps are identifiable by diffuse and increased β-catenin staining confirming that the cells they contain had undergone LOH. In contrast, in *Apc*^*Min/+*^ heterozygous tissue, β-catenin localized to the lateral membranes, and only these cells exhibited strong nuclear pSTAT3 after IL-22 injection. These data confirm that epithelial cells heterozygous for APC mutation respond to IL-22 similarly to WT cells, whereas homozygous mutant *Apc*^*Min/Min*^ cells do not respond to IL-22 *in vivo*.

**Figure 6.**
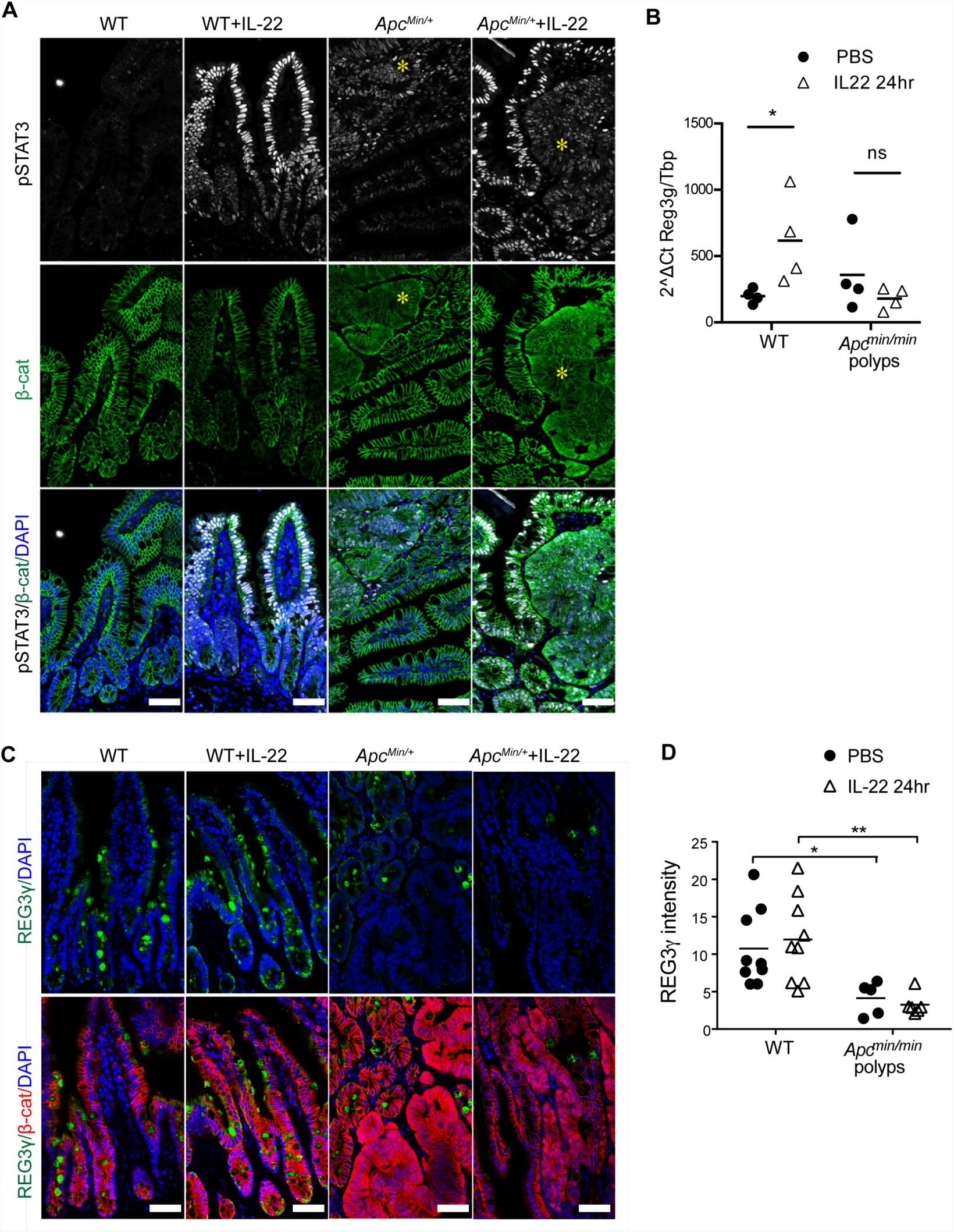
Reduced IL-22 responses in cells in *Apc*^*Min/Min*^ small intestinal polyps *in vivo*. Mice aged between 88-95 days were injected i.p. with 1µg IL-22 or PBS. (A) Small intestine was harvested after 1 hour and tissue stained with antibodies against pSTAT3 (Tyr 705) and β-catenin. Nuclei were stained with DAPI. Asterisk indicates polyps. (B) Small intestine was harvested 24 hours after IL-22 injection. RNA was isolated and RT-qPCR was performed. Data shows the level of *Reg3g* mRNA relative to *Tbp*. *P<0.05, two-way ANOVA. (C) Small intestine was harvested 24 hours after IL-22 injection and IHC was performed using antibodies against RegIIIγ and β-catenin. Nuclei were stained with DAPI. 3 independent experiments were performed. Number of mice used in total: WT=3; *Apc*^*Min/+*^ =6. Scale bar = 50µm. (D). Mean fluorescence intensity for RegIIIγ in the epithelial area analysed using Fuji software. Each dot represents the mean fluorescence intensity in 1 confocal image. *P<0.05, **P<0.01, t-test.

We next asked whether the lack of pSTAT3 induction resulted in lower STAT3 target gene expression in *Apc*^*Min/Min*^ polyps. *Reg3g* was one of the most highly induced antimicrobial genes in response to IL-22 in organoids. Consistent with our *in vitro* data, *Reg3g* mRNA was induced in response to IL-22 in WT, but not in *Apc*^*Min/Min*^ tissue (Fig. 6B). We also observed significantly lower expression of RegIIIγ peptides in polyps compared to WT or *Apc*^*Min/+*^ epithelia with or without IL-22 injection (Fig. 6C & 6D). Thus the loss of IL-22 responsiveness in homozygous *Apc*^*Min/Min*^ cells was also recapitulated *in vivo*.

### IL-22 induces oxidative stress and promotes early intestinal tumorigenesis

Since transformed *Apc*^*Min/Min*^ tissue responded poorly to IL-22, we reasoned that IL-22 must contribute to intestinal tumorigenesis by acting on *Apc*^*Min/+*^ cells, before LOH drives loss of IL-22 responses. Many inflammatory cytokines induce production of reactive oxygen and nitrogen species as part of the defence response. Importantly, these free radicals can also cause DNA damage. We hypothesized that IL-22–induced oxidative stress could promote tumorigenesis by driving DNA damage. Our RNAseq data indicated that IL-22 up-regulated the inducible Nitric oxide synthase (iNOS, *Nos2*) and Dual oxidases (*Duox1/2) (*Fig. 1C, 2C), as well as several genes involved in DNA repair (e.g. *Parp9, Gadd45g*). We confirmed that IL-22 induced iNOS protein expression in both WT and heterozygous *Apc*^*Min/+*^ cells (Fig. 7A, B). Moreover, the DNA damage marker γH2AX was increased by IL22 in both WT and *Apc*^*Min/+*^ cells (Fig. 7A, B). This suggested that IL-22–induced DNA damage could contribute to loss of heterozygosity and initiate APC mutation-dependent intestinal transformation.

**Figure 7.**
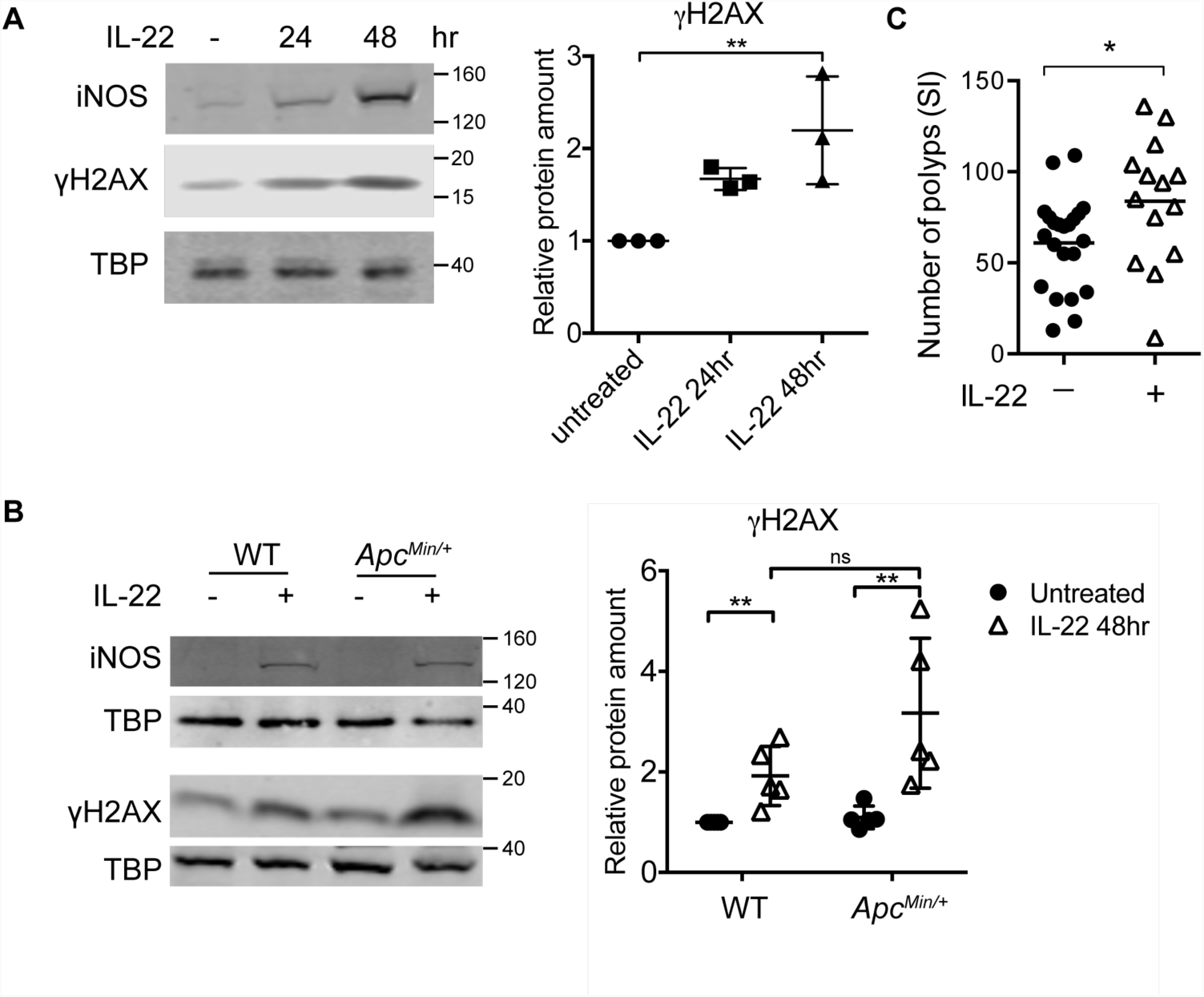
IL-22 increases iNOS and DNA damage in *Apc*^*Min/+*^ organoids. (A). Western blot of iNOS or γH2AX in WT organoids treated with IL-22 (10ng/ml) for 24 or 48 hours. (B). Western blotting for iNOS or γH2AX in WT or *Apc*^*Min/+*^ organoids treated with IL-22 (10ng/ml) for 48 hours. TBP was used as loading control. The ratio of γH2AX to TBP in each sample was normalised to untreated control in each experiment. At least 3 independent experiments were performed. *P<0.05, paired t-test. (C) *Apc*^*Min/+*^ mice aged 27-30 days old were injected twice a week for 4 weeks with 1µg IL-22 or PBS, or were left untreated. Untreated and PBS injected mice were pooled together for the ‘control’ cohort (there was no difference in the tumour number in these two cohorts). * P<0.05, unpaired t-test.

To test this idea, we injected young (4-5 week old) *Apc*^*Min/+*^ mice with low levels (1µg) of IL-22 twice a week for 4 weeks, to mimic underlying inflammation. Indeed, after 5 weeks we found increased numbers of polyps/tumours in the SI of *Apc*^*Min/+*^ mice that had been injected with IL-22, than in animals that had not been injected with IL-22 (Fig. 7C). These observations support the idea that IL-22 could drive tumorigenesis by promoting LOH in *Apc*^*Min/+*^ cells in mice.

## Discussion

The cytokine IL-22 has emerged as an important player in intestinal tumorigenesis. IL-22 can increase cell proliferation (Lindemans *et al*, 2015) and enhance cancer stemness by increasing expression of core stem cell genes (Kryczek *et al*, 2014). We investigated the mechanism by which IL-22 contributes to tumorigenesis in the *Apc*^*Min/+*^ mouse model for cancer. Unexpectedly, we found that transformed *Apc*^*Min/Min*^ cells in both organoids and in animals responded poorly to IL-22. We further showed that this was likely due to low IL-22 receptor expression and reduced STAT3 transcriptional activity in *Apc*^*Min/Min*^ cells. Our data suggest that rather than contributing to intestinal tumorigenesis by augmenting the proliferation of transformed cells, IL-22 acts at an early stage to drive genetic instability in *Apc*^*Min/+*^ cells to increase the occurrence of transforming LOH events.

A significant prediction of our work is that the absence of IL-22 responses in *Apc*^*Min/Min*^ cells may lead to increased intestinal inflammation due to reduced immune defence of the intestinal barrier. Previous reports and our own transcriptome analysis show that the main function of IL-22 is to up-regulate defence and immune responses (Zheng *et al*, 2008; Behnsen *et al*, 2014), through upregulation of mucus production, mucus glycosylation and antimicrobial peptide production. IL-22 producing ILCs can prevent bacterial translocation across the intestinal barrier. Correspondingly, lack of IL-22, or IL-22 receptor contributes to a heightened inflammatory milieu in the intestinal epithelium, and increased sensitivity to bacterial infection due to increased systemic dissemination of bacteria (Vaishnava *et al*, 2012; Loonen *et al*, 2014; Pham *et al*, 2014). In this context, previous studies showed compromised intestinal barrier integrity and increased inflammation in *Apc*^*Min/+*^ and *Apc*^*Floxed/wt*^; *Cdx2-Cre* mice (Li *et al*, 2012; Grivennikov *et al*, 2012). Consistent with the lack of IL-22 responses in *Apc*^*Min/Min*^ cells, we found that the antimicrobial peptide RegIIIγ was nearly undetectable in polyps in *Apc*^*Min/+*^ mice. Lack of RegIIIγ can result in increased bacterial translocation across the intestinal epithelial barrier, and the decreased mucosal protection in *Reg3g*^-/-^ mice correlates with increased inflammation (Loonen *et al*, 2014; Vaishnava *et al*, 2012). Thus, the lack of RegIIIγ expression in polyps, even in the presence of IL-22, may cause impaired barrier functions and increased inflammation in *Apc*^*Min/Min*^ polyps.

In the absence of IL-22 responses in transformed *Apc*^*Min/Min*^ cells, the most likely explanation for the tumour-promoting effect of IL-22 in *Apc*^*Min/+*^ mice is provided by our discovery that IL-22 induces expression of iNOS and Duox1/2. This is predicted to result in increased RNS and ROS production and DNA damage. Consistent with the idea that RNS could accelerate polyp formation, inhibition of nitric oxide production reduced tumour loads in *Apc*^*Min/+*^ mice (Ahn & Ohshima, 2001). In addition, studies in colitis-associated carcinogenesis mouse model using *H. hepaticus* infection, showed that IL-22 could induce DNA damage via nitric oxide-dependent mechanism to promote dysplasia (Wang *et al*, 2017). Infection of 4 week old *Apc*^*Min/+*^ mice with *Citrobacter rodentium*, a colonic attaching and effacing pathogen that induces a potent IL-22 response, increased colonic tumour burden (Newman *et al*, 2001). We also observed increased small intestinal tumour burden upon multiple IL-22 injections into young preneoplastic *Apc*^*Min/+*^ mice. Together, these data support the idea that IL-22 has mutagenic effects on non-transformed cells, mediated by iNOS. This in turn could explain the reduction in tumour numbers in IL22^-/-^ *Apc*^*Min/+*^ compared to IL22^+/+^ *Apc*^*Min/+*^ mice.

It is intriguing that APC-mutant cells selectively downregulate IL-22–dependent pSTAT3 signalling, whereas IL-6 induced pSTAT3 is unaffected. One explanation for the lower IL-22R expression could be reduced stability of the protein. The IL-22R is phosphorylated by GSK3, which prevents its proteasomal degradation (Weathington *et al*, 2014). In the presence of mutant APC, GSK3 activity is reduced (Valvezan *et al*, 2014), which may partially explain the low surface expression of the IL-22 receptor in *Apc*^*Min/Min*^ cells. However, it should be noted that our RNAseq data indicated differential expression of nearly 5,000 genes in *Apc*^*Min/Min*^ compared to WT organoids, suggesting that there are other possible reasons for the loss of IL-22 responsiveness in APC-mutant cells.

Mutations in APC are found in up to 80% of non-hereditary CRC (Kinzler & Vogelstein, 1996). However, we are not suggesting that all transformed epithelial cells cannot respond to IL-22. When analysing a panel of human cell lines, we found that a subset of them did not respond to IL-22, and some of these had reduced expression of the IL-22 receptor ((Brand *et al*, 2006) and data not shown). On the other hand, high levels of IL-22 have been found in many CRC samples, and is associated with CRC development (Jiang *et al*, 2013; Kirchberger *et al*, 2013; Huang *et al*, 2015). Whether the presence of IL-22 is important at an early stage for the development of tumours, or for maintaining tumour progression is not yet clearly established (Hernandez *et al*, 2018). Importantly, our data advocates exercising caution when considering IL-22 as a therapeutic target for treatment of established CRC, and suggest that patients need to be stratified based on expression of the IL-22 receptor, STAT3 repressors and possibly other mutations acquired by cancerous cells.

## Supporting information

## Acknowledgements

This work was supported by the National Centre for the Replacement, Refinement & Reduction of Animals in Research (To ISN and MS, NC/M001156/1) and by the Wellcome Trust and Royal Society (Sir Henry Dale Fellowship to MS, 206246/Z/17/Z). We are grateful to Prof. Lora Hooper and her lab (University of Texas Southwestern Medical Center, Texas, USA) for providing RegIIIγ antibodies and advice. We thank Dr Sarah Thompson and Don Tennant for assistance with mouse models (Biological Resource Unit, University of Dundee), Dr. Rosemary Clarke (Flow Cytometry Facility, University of Dundee) for assistance with flow cytometry and Dr. Graeme Ball (Dundee Imaging Facility, University of Dundee) for assistance with image analysis.

## Author contributions

YC, ISN and MS conceptualized and designed the study, Y.C. performed most experiments and analysed results, MS performed experiments and analysed data, IPN and MV assisted in *in vivo* experiments and data collection, IM provided key reagents and conceptual help, YC, ISN and MS interpreted data, MS and YC wrote the manuscript with input from all co-authors.

## Materials and methods

### Mice and treatment

All experiments involving animals were performed under the UK Home Office guidelines and were approved by the Home office Licensing committee (Project licenses PPL60/4172, 70/8813, PD4D8EFEF). Healthy C57BL/6J wild type male or female adult mice between 60-120 days old bred in-house were used for preparing WT organoids. For *APC*^*Min/Min*^ organoids or *APC*^*Min/+*^ organoids, *APC*^*Min/+*^ male or female adult mice age approximately at 90-day-old or 60-day-old were used respectively. Mice were maintained in a standard barrier facility on a 12hour light/dark cycle at 21°C, 55-65% relative humidity. Mice were tested negative for all pathogens on the current FELASA list, except for *Helicobacter spp.* Mice were maintained in standard individually ventilated cages with Eco pure chips 6 and fed an autoclaved R&M3 diet (Special Diet Services) and autoclaved water *ad libitum*, and cages were changed at least every two weeks.

For short-term IL-22 treatment, WT or *Apc*^*Min/+*^ mice aged approximately 90 days old were injected intraperitoneally with 1µg IL-22 and PBS-injected mice were used as a non-stimulated control. Mice were sampled 1-24 hours after treatment. For longer term treatments, WT or *Apc*^*Min/+*^ mice aged 28-35 days old were injected intraperitoneally with 1µg of IL-22 or PBS or left untreated, twice every week for 4 weeks. 1 week after the last injection, intestines were harvested and polyp load assessed.

### Intestinal organoid culture and treatment

Small intestine of CL57BL/6J was washed with PBS and was longitudinally opened. Villi were moved by scraping with a glass coverslip. Following further PBS washes, small intestine was incubated in 3mM EDTA in PBS at 4°C for 20 minutes. After being shaken for 1 minute, small intestine was removed and the crypts were collected by centrifugation at 600rpm at 4°C. Crypts were washed with PBS and incubated with TrypLE Express (GIBCO) at 37 °C for 5 minutes. Cells were suspended in Advanced DMEM/F12 (ADF) medium (GIBCO) and were passed through a 40µm cell strainer (Greiner). After centrifugation at 2000rpm for 3 min, cells were re-suspended in Matrigel (Corning) and were plated into 24-well plate. For WT or *Apc*^*Min/+*^ organoid culture, cells were incubated in a 37°C 5% CO_2_ incubator for 15 min and were then supplemented with crypt medium (ADF medium supplemented with HEPES (GIBCO), GlutaMax-1 (GIBCO), N-acetylcysteine (10mM, Sigma), N2 (GIBCO) and B27 (GIBCO), 100U/ml penicillin/ 100µg/ml streptomycin (Invitrogen) containing EGF (50ng/ml, GIBCO), Noggin (100ng/ml Peprotech), R-Spondin conditioned-medium, valproic acid (1µM, Sigma) and ROCK Inhibitor Y27632 (Sigma)). After 48-hour incubation, medium was replaced by crypt medium supplemented with EGF, Noggin and R-Spondin. For *Apc*^*Min/Min*^ organoid culture, crypt medium containing EGF, Noggin and R-Spondin was used for the first 48 hours. *Apc*^*Min/Min*^ organoids were then cultured in crypt medium alone for at a week before replaced by crypt medium supplied with EGF, Noggin and R-Spondin. Medium was replaced every 2 days. Organoids were passaged every 4∼6 days. Organoids were used for experiments at day 3 after passaging. For IL-22 stimulation experiment, organoids were treated with IL-22 (Peprotech). Concentration and duration are described in the figure legends. For HDAC inhibition, organoids were incubated with 1mM sodium butyrate (NaBu) (Sigma) for 16 hours before stimulated with IL-22 (10ng/ml) along with media containing 1mM NaBu for 3 hours.

### Immunohistochemistry (IHC) staining

For IHC staining, small intestine was removed and fixed in 4% PFA at 4°C overnight. Followed by processing and paraffin embedding, specimens were cut into 5µm sections and placed onto SuperFrost Plus™ Adhesion Slides (Thermo Scientific) and dried at 60°C overnight. Slides were de-paraffined with Histo-Clear (National Diagnostics) and dehydrated through graded ethanol solutions. For antigen retrieval, EDTA buffer (1mM EDTA, 0.05% Tween 20, pH 8) was used for pSTAT3 and β-catenin co-staining, and Sodium Citrate buffer (10mM Sodium Citrate, 0.05% Tween 20, pH 6.0) was for RegIIIγ and β-catenin co-staining. Slides were rinsed in PBS and incubated with blocking buffer 1 (PBS, 2%BSA) for 15 minutes, followed by incubation with blocking buffer 2 (PBS, 1%BSA, 0.3% Triton) for 15 minutes. Slides were immersed with primary antibodies diluted in blocking buffer 2 for 3 hours. Primary antibodies used include anti-β-catenin (1:200, BD, 610154), pSTAT3 (Tyr705) (1:200, Cell Signaling, 9145) and anti-RegIIIγ (1:200, gift from Lora Hooper). After TBST (TBS, 0.1% Tween-20) washes, slides were incubated with Alexa Fluor™ fluorophore conjugated secondary antibody (1:500, Invitrogen) and DAPI (1µg/ml, Invitrogen) for 30 minutes. After TBST washes, the cover slides were mounted on glass slides with Prolong Gold Antifade media. Images were analysed in Imaris software (Bitplane) or Fuji software (Schindelin *et al*, 2012).

### Immunofluorescence staining of intestinal organoids

For imaging and immunofluorescence, organoids were grown in Matrigel on 8-well µ-slides (Ibidi). Organoids were fixed in pre-warmed (37°C) 4% paraformaldehyde (pH7.4, Sigma) for 20 minutes at 37°C. Organoids were permeabilized by using permeabilization buffer (PBS, 1% Triton X-100) for 1 hour, followed by blocking step with blocking buffer (PBS, 1% BSA, 3% normal goat serum, 0.2% Triton X-100) for 1 hour at room temperature. Organoids were then incubated with primary antibody diluted in working buffer (PBS, 0.1% BSA, 0.3% normal goat serum, 0.2% Triton X-100) overnight at room temperature. Primary antibodies used were as follows: anti-Phospho-Stat3 (Tyr705) (1:100, Cell Signaling, 9131). After washed with working buffer, organoids were incubated with working buffer with Alexa Fluor™ fluorophore conjugated secondary antibody (1:500, Invitrogen) and Hoechst 33342 (5µg/ml, Invitrogen) overnight at room temperature. Organoids were then washed with working buffer and were mounted with ProLong^®^ Gold antifade mountant (Invitrogen). Immunofluorescence was visualized under Zeiss 710 confocal microscope. Images were analysed by using Imaris software (Bitplane).

### Western blotting

Organoids were washed with ice-cold PBS and cell pellets were lysed in lysis buffer (40mM Tris-HCl pH7.5, 120mM NaCl, 0.27M sucrose, 1mM EDTA pH8, 1mM EGTA, 10mM β-glycerophosphate, 5mM Sodium pyrophosphate, 1% Triton X-100) containing protein and phosphatase inhibitors. Proteins are separated on a NuPAGE™ 4-12% Bis-Tris Protein Gels (Invitrogen) and are transferred to Protran nitrocellulose membrane (0.2 µm, GE Healthcare). Immunoblotting was performed with primary antibodies diluted in 5%BSA/TBS-Triton overnight at 4°C. Primary antibodies used were as follows: anti-Phospho-Stat3 (Tyr705) (1:2000, Cell Signaling, 9131); anti-Stat3 (1:2000, Cell Signaling, 9139); anti-Phospho-Stat3 (Ser727) (1:1000, Cell Signaling, 9134); anti-Phospho-Stat1(Tyr 701) (1:1000, Cell Signaling, 9167); anti-Stat1 (1:2000, BD, 610185); anti-TBP (1:2000, Proteintech, 66166-1-lg), anti-HDAC1 (1:1000, 06-720, Millipore), anti-HDAC2 (1:1000, Cell Signaling, 5113), anti-iNOS (1:200, Abcam, ab15323), anti-γH2AX (1:500, Biolegend, 613401). Blots were incubated with IRDye800/700-conjugated secondary antibodies (Rockland, 1:5000) for 1 hour at room temperature. Proteins were detected with LiCor Odyssey Imager (Licor Biosciences). Western blots were analysed in Image Studio Lite 5.2 software (Licor Biosciences).

### Quantitative real-time reverse transcription polymerase chain reaction (RT-qPCR)

Total RNA was isolated from organoids by using NucleoSpin RNA isolationKit (Macherey-Nagel). cDNA was prepared from total RNA by using qScript cDNA synthesis kit (Quanta). Real-time PCR reaction was performed by using PerfeCTa SYBR green FastMix for iQ (Quanta) in a qPCR machine (Bio-Rad CFX-Connect Real-Time system). Primers (Eurofins MWG) used are shown in supplementary table 1.

### RNA sequencing (RNAseq)

WT or *Apc*^*Min/Min*^ organoids were stimulated with IL-22 (2ng/ml) for 3 hours. Total RNA was isolated from organoids by using NucleoSpin RNA isolation Kit (Macherey-Nagel). RNAsep was performed and analysed by Turku Centre for Biotechnology, University of Turku and Åbo Akademi University. In brief, RNA was prepared for the sequencing using Illumina TruSeq Stranded mRNA sample Preparation Kit. mRNA was sequenced with the HiSeq 2500 instrument using single-end sequencing chemistry and 50bp red length. STAR version 2.5.2b was used to align the reads against the reference genome. *Subreads* package (v. 1.5.1) was then used to be associated with known genes and to count the number of reads associated with each gene.

For group comparison, statistical testing between the groups was performed in R package Limma (Bioconductor) (Smyth *et al*, 2005). For functional analysis, topGO and GOstats packages in R/Bioconductor were used for the enrichment analysis of differentially expressed filtered gene lists. Further analyses were performed using the DAVID Functional Annotation tools. Heatmaps were prepared using the Morpheus software (https://software.broadinstitute.org/morpheus), and the Viridis color palette.

### Flow cytometry

To prepare single cell suspension, organoids were dissociated by incubation with TrypLE Express at 37°C for 5 minutes. Cells were collected by centrifugation at 2000 rpm for 2 minutes. For IL22RA1 staining, cells were stained with IL22RA1 antibody (1:100, Proteintech, 13462-1-AP) for 30 minutes on ice. After washed with staining buffer (PBS, 1%FBS, 2mM EDTA), cells were incubated with Donkey Anti-Rabbit Alexa Fluor 647 antibody (1:500, ThermoFisher) for 30 minutes on ice. After washed with staining buffer, cells were re-suspended in staining buffer containing DAPI (2.5µg/ml, Invitrogen).

For STAT3 staining, after incubation with TrypLE, single cells were incubated in 0.5 ml PBS containing LIVE/DEAD fixable blue dead cell stain kit (1:1000, Invitrogen) on ice for 10 minutes. Cell were washed with staining buffer and then collected by centrifugation at 2000 rpm for 2 minutes. Cells were fixed with 3ml pre-warmed 0.5% PFA/PBS at 37°C for 15 minutes and then washed with staining buffer. Permeabilization was performed by incubation with ice-cold 70% ethanol on ice for 10 minutes. After wash with staining buffer, cells were stained with STAT3 antibody (1: 200, Cell Signaling, 9139) for 1 hour. After washed with staining buffer, cells were stained with goat anti-mouse Alexa 568 antibody (Invitrogen) for 1 hour. After wash with staining buffer, cells were re-suspended in 500ul staining buffer and analysed by LSRFortessa Cell Analyzer (BD). Flow cytometry data was analysed by using FlowJo software V9 (FlowJo, LLC).

### Statistical analysis

Statistical analysis was performed by using Prism 5.0b (GraphPad).

