## Supplementary material for "Loss of Adenomatous polyposis coli function renders intestinal epithelial cells resistant to the cytokine IL-22"

(Supplemental data)

Yu Chen[1,2], Ian P. Newton[1], Maud Vandereyken[2], Ignacio Moraga[3], Inke Näthke[1*], Mahima Swamy[2*]

[1]Cell and Developmental Biology, [2]MRC Protein Phosphorylation and Ubiquitylation Unit (PPU), [3]Cell Signalling and Immunology, University of Dundee, Dundee,

DD1 5EH, UK

**A**

| Comparison | FC | p-value | Total | Up | Down |
|---|---|---|---|---|---|
| WT+IL-22 vs. WT | 2.0 | 0.05 | 437 | 263 | 174 |
| $Apc^{Min/Min}$ vs. WT | 2.0 | 0.05 | 4881 | 2517 | 2364 |
| $Apc^{Min/Min}$+IL-22 vs. $Apc^{Min}$ | 2.0 | **0.10** | 12 | 11 | 1 |
| $Apc^{Min/Min}$+IL-22 vs. WT+IL-22 | 2.0 | 0.05 | 4674 | 2381 | 2293 |

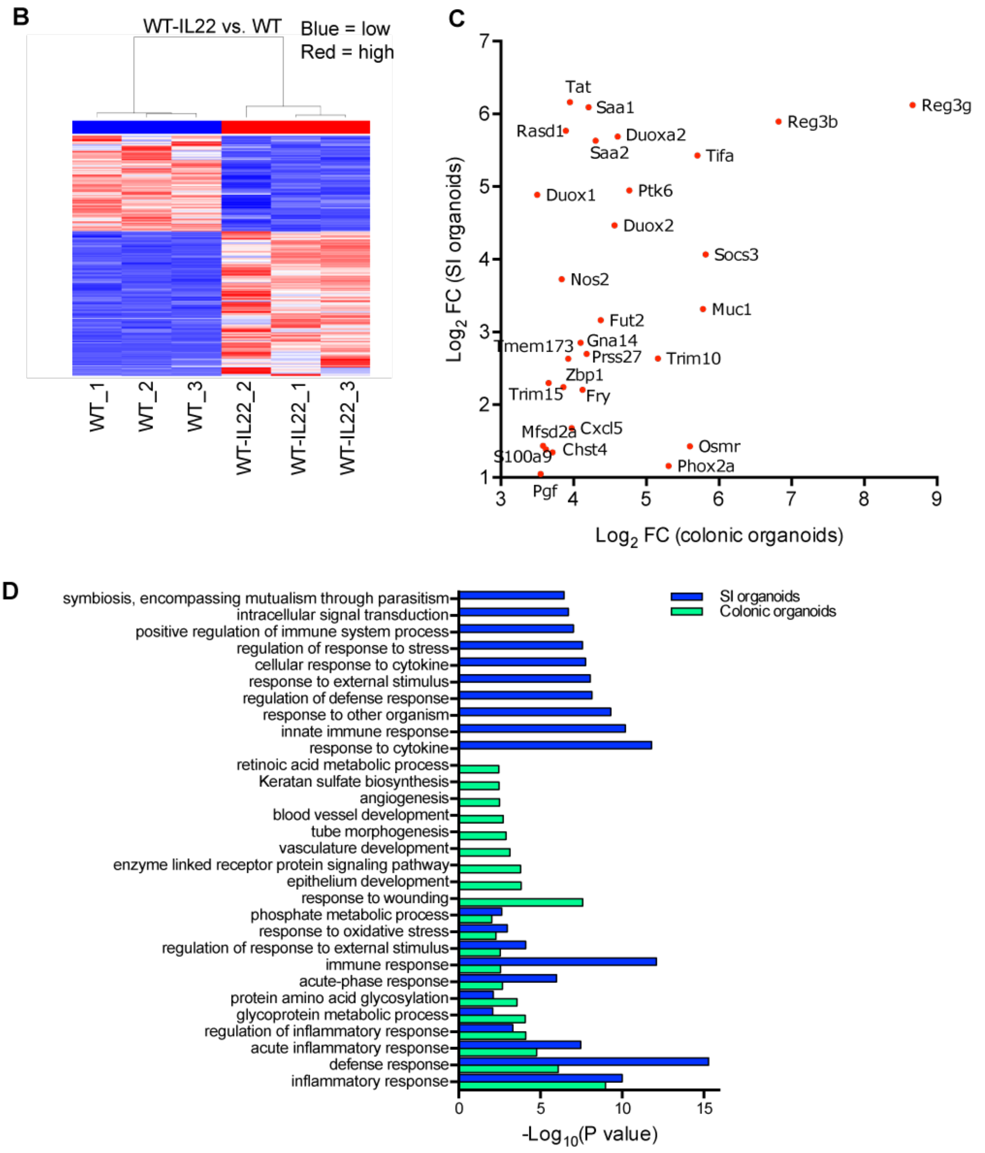


**Supplementary figure 1. Transcriptome comparison of IL-22 responses in WT and *Apc* *Min/Min* organoids**

(A) Table shows the number of genes significantly regulated by IL-22 in WT or *Apc*$^{Min/Min}$ organoids, and the thresholds used to filter the data. FC: fold change. (B) Heat map plot for differentially expressed genes between WT organoids treated with or without IL-22 for 3 h. Heatmap: red=higher expression, blue=lower expression, relative to the row mean expression of the gene across all samples. (C) Out of the top 50 IL-22 upregulated genes in colonic organoids (Pham *et al*, 2014), only 29 had a fold change>2 in IL-22 treated small intestinal organoids. (D) The top 20 biological processes (filtered, GO_BP_FAT) regulated by IL-22 in colonic and small intestinal organoids.

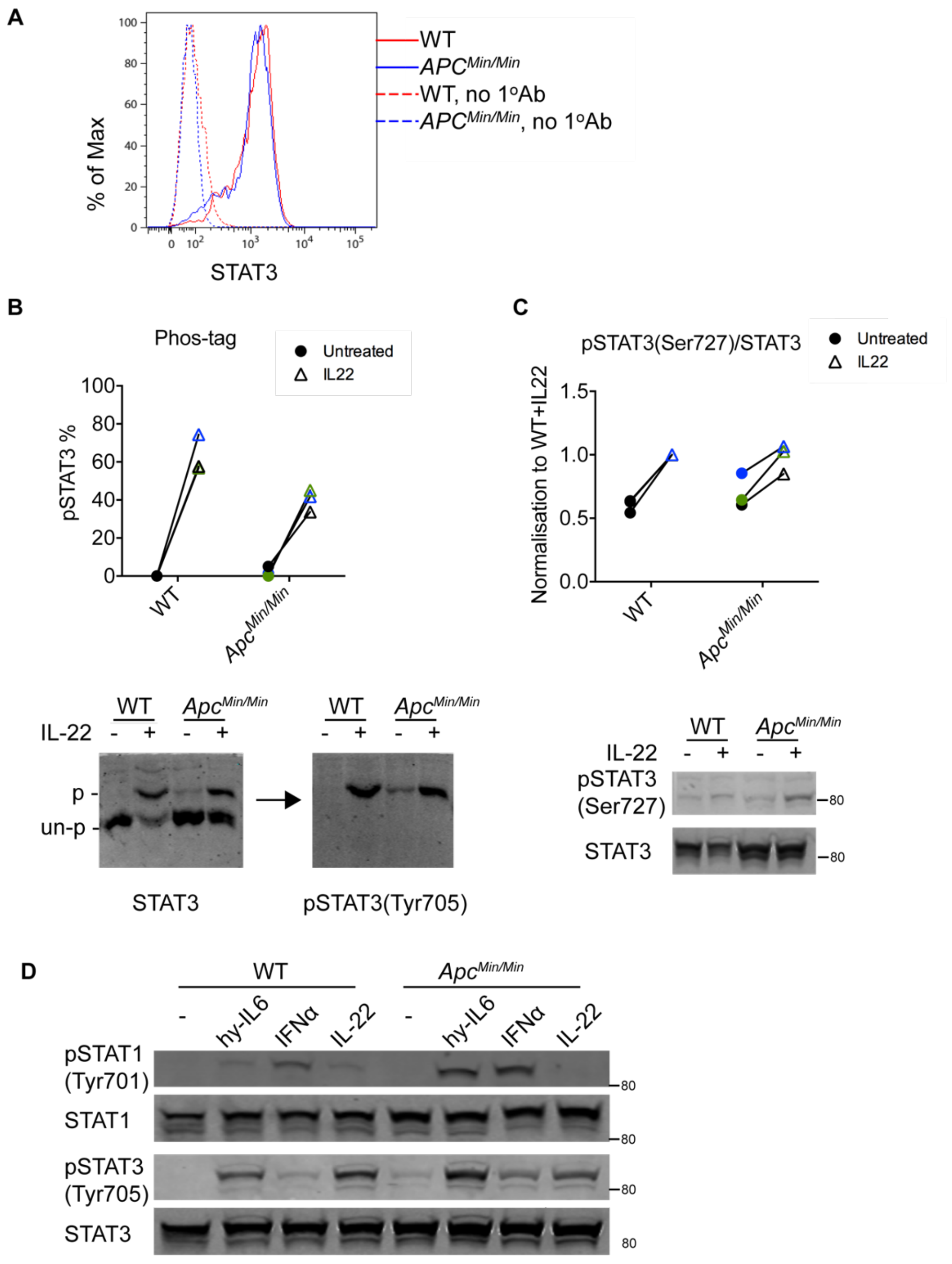


**Supplementary figure 2. IL-22 increases pSTAT3 (Ser727)**

(A) Flow cytometry analysis of STAT3 expression in WT and *Apc*$^{Min/Min}$ organoids. (B) Phos-tag gels were used to separate phosphorylated and non-phosphorylated STAT3. Immunoblot for STAT3 shows non-phosphorylated (lower band) and phosphorylated (upper band) STAT3 protein. The same membrane was incubated with anti-pSTAT3 (Tyr705) to confirm the identity of the upper band as pSTAT3. Plot shows the percentage of total STAT3 that is phosphorylated.

(C) Western blot analysis shows pSTAT3 (Ser727) levels in WT and *Apc*$^{Min/Min}$ organoids with or without IL-22 stimulation (10ng/ml) for 0.5 hour. Data show the ratio of pSTAT3(Ser727) to total STAT3 in each sample normalised to that in WT organoids treated with IL-22 in each experiment. (D) Western blot of pSTAT3(Tyr705), STAT3, pSTAT1(Tyr701) or STAT1 in WT and *Apc*$^{Min/Min}$ organoids treated with IL-22 (10ng/ml), hyper IL-6 (hy IL-6, 50μM) or IFNα (1000U/ml) for 0.5 hours.

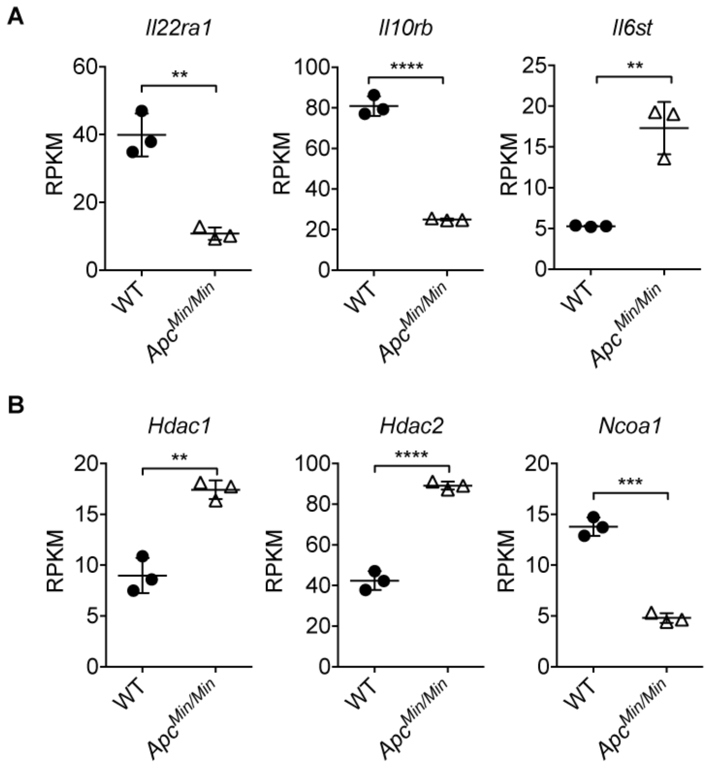


**Supplementary figure 3. *Apc* $^{Min/Min}$ organoids express lower mRNA levels of IL-22 receptor genes**

RNAseq data for mRNA levels of (A) *Il22ra1*, *Il10rb, Il6st* and (B) *Hdac1*, *Hdac2* and *Ncoa1* in WT and *Apc*$^{Min/Min}$ organoids. RPKM: reads per kilo base per million mapped reads. **P<0.01, ***P<0.001, and ****P<0.0001, t-test.

**Supplementary Table 1**. Primers Sequences used for qPCR

| Gene | Forward primer | Reverse primer |
|---|---|---|
| *Adar* | ggaagaagactcggagaaacc | tcccagagaacaaggatgttg |
| *Duox2* | gcactgtgcagaacagctaggacaac | acctcatcaccttcttgcgggag |
| *Fut2* | agagccaagctggaagatacac | atcaaggtggcgtctctctg |
| *Lcn2* | ccatctatgagctacaagagaacaat | tctgatccagtagcgacagc |
| *Lrg1* | ccatgtcagtgtgcagattc | aagagtgagaggtggaagag |
| *Nos2* | ttcagcacatctgcagacac | agcctgaagtcatgtttgcc |
| *Ptk6* | gccgtgcgacattacaggat | taggccatgagacaggctct |
| *Reg3g* | accatcaccatcatgtcctg | ggcatctttcttggcaactt |
| *Saa1/2* | ctgcctgccaaatactgagagtc | ccacttccaagttcctgtttattac |
| *Saa3* | gctggcctgcctaaaagatactg | gcatttcacaagtatttattcagc |
| *Socs3* | gcgggcacctttcttatcc | tccccgactgggtcttgac |
| *Tbp* | ggggagctgtgatgtgaagt | ccaggaaataattctggctcat |
| *Tifa* | acgcaattccaacatgtgcc | ccgagctcctggttgtctac |
| *Usp18* | ttgggctcctgaggaaacc | cgatgttgtgtaaaccaaccaga |
